# Conservation management strategy impacts inbreeding and mutation load in scimitar-horned oryx

**DOI:** 10.1101/2022.06.19.496717

**Authors:** Emily Humble, Martin A Stoffel, Kara Dicks, Alex D Ball, Rebecca M Gooley, Justin Chuven, Ricardo Pusey, Mohammed Al Remeithi, Klaus-Peter Koepfli, Budhan Pukazhenthi, Helen Senn, Rob Ogden

## Abstract

In an age of habitat loss and overexploitation, small populations, both captive and wild, are increasingly facing the effects of isolation and inbreeding. Genetic management has therefore become a vital tool for ensuring population viability. However, little is known about how the type and intensity of intervention shape the genomic landscape of inbreeding and mutation load. We address this using whole genome sequence data of scimitar-horned oryx (*Oryx dammah*), an iconic antelope that has been subject to contrasting management strategies since it was declared extinct in the wild. We show that unmanaged populations are enriched for long runs of homozygosity (ROH) and have significantly higher inbreeding coefficients than managed populations. Additionally, despite the total number of deleterious alleles being similar across management strategies, the burden of homozygous deleterious genotypes was consistently higher in unmanaged groups. These findings emphasise the risks associated with deleterious mutations through multiple generations of inbreeding. As wildlife management strategies continue to diversify, our study reinforces the importance of maintaining genome-wide variation in vulnerable populations and has direct implications for one of the largest reintroduction attempts in the world.

**Significance statement:** Conservation genetic management is becoming increasingly important for safeguarding and restoring wildlife populations. Understanding how the intensity of intervention influences genomic components of fitness is therefore essential for supporting species viability. We investigate the impact of contrasting management strategies on the genomic landscape of inbreeding and mutation load in captive populations of scimitar-horned oryx. We reveal how several decades of management have prevented the formation of long runs of homozygosity and masked the expression of deleterious mutations. Our findings highlight the dynamics between inbreeding, mutation load and population size and have direct implications for future management of threatened species.

## Main text Introduction

Captive populations have become an essential insurance against extinctions in the wild (1). However, due to inbreeding and drift, they are intrinsically vulnerable to reduced genetic variation and the expression of partially recessive deleterious mutations (2–6). It is therefore of paramount importance that appropriate plans are in place to safeguard their potential as source populations. *Ex-situ* management strategies fall along a continuum from high-intensity pedigree-based breeding (7), to low-intensity pedigree-free group management (8, 9), to a complete absence of breeding intervention whatsoever. Empirical evidence on how these approaches influence the combined landscape of inbreeding and deleterious variation is limited (10, 11). As wildlife management strategies begin to diversify (12–15), there is a pressing need to leverage current genomic techniques to understand how genetic components of fitness are impacted by conservation intervention.

Alongside this, recent debate on the significance of neutral genetic variation in conservation biology has raised practical considerations for sourcing populations for restorations (16–20). For example, an increasing number of studies are uncovering genomic evidence for purging in the wild (21–26), some of which have used this to challenge the small population paradigm of conservation biology (27–29). Furthermore, simulation-based studies on the interaction between effective population size, genetic variation and extinction risk have called for more emphasis on functional genomic variation in genetic rescue attempts (18, 19). These observations go against decades of empirical and theoretical work in favour of maximising genetic variation to enhance population viability (30–33) including recent studies highlighting the complex dynamics of deleterious mutation frequencies in small populations (34–38). Founder selection for translocations rests on a complex set of considerations, with genetics making up only one component (39). In most cases, conservation practitioners will favour a unifying strategy to minimise risk and maximise return (40–42). In light of this, empirical data on the patterns of inbreeding and deleterious mutations in species undergoing active conservation management are urgently required.

*Ex-situ* populations of scimitar-horned oryx provide an excellent opportunity to evaluate the genomic consequences of management in the context of a global reintroduction. This iconic antelope was once widespread across North Africa, yet during the 20^th^ century, hunting and land-use competition led to their rapid population decline and eventual extinction from the wild (43). Prior to disappearing, captive populations had already been established from what is thought to be less than 100 animals originating from Chad in the 1960s (43). In the following years, the *ex-situ* population has grown to reach approximately 15,000 individuals (44). Around 1,000 of these are held within coordinated breeding programmes, but the vast majority are held in collections in places like Texas and the Arabian Peninsula where little to no genetic management takes place. Crucially, the scimitar-horned oryx is now being reintroduced back into its former range and *ex-situ* populations with varying management strategies have been used to source individuals for release. Here, we use runs of homozygosity (ROH) and predicted deleterious mutations to evaluate the impacts of captive-breeding practices on inbreeding and mutation load in scimitar-horned oryx, and discuss the implications for its ongoing management.

## Results

We generated whole-genome sequence data for 49 scimitar-horned oryx from four *ex-situ* populations. Two of these, the EAZA *Ex situ* Programmes (EEP, *n* = 8) and the USA (*n* = 17), represent captive populations where genetic management practices are in place. The EEP population comprised individuals from zoological institutions across Europe. The USA population comprised individuals from both privately owned ranches and institutions within the AZA Species Survival Plan ® (SSP). The remaining populations from the Environment Agency – Abu Dhabi originate from two genetically unmanaged collections in the United Arab Emirates (EAD A: *n* = 9 and EAD B: *n* = 15). Census sizes for the EEP and SSP population are approximately 619 and 223, respectively, while those for EAD A and EAD B are approximately 3,000 and 70. For further details on population origins, management strategies and sampling approach, please refer to the Supplementary Material.

High coverage sequencing (∼15X) was performed for 20 of the individuals and the remaining 29 were sequenced at a lower depth (6–8X, Table S1). Sequencing reads were mapped to the scimitar-horned oryx reference genome (45) and to account for coverage biases, SNPs and genotype likelihoods were called after downsampling high coverage individuals (see Methods for details). Analysis of population structure using NGSadmix and PCAngsd detected differentiation between the four sampling groups (Figures S1–3). Individual admixture proportions highlighted two major ancestral source populations (Figures S1A), with further hierarchical structure being resolved up to values of K=4 (Figures S1B and S2), corresponding to the four *ex-situ* groups. PCA distinguished EEP and USA populations as discrete clusters along PC2 and PC3, while EAD A and EAD B clustered separately along PC1 (Figure S4).

### Levels of inbreeding across management strategies

To investigate how genomic patterns of inbreeding vary with management strategy, we examined the ROH landscape across individuals (Figure 1). The average number and total length of ROH was 247 (min = 65, max = 638) and 2.0 Mb (0.5–22.0 Mb) respectively, which on average spanned 20% of the autosomal genome (min = 0.03, max = 0.55, Figure 1A and Figure S4). Oryx from managed populations had significantly lower inbreeding coefficients (*F*_*ROH*_) than oryx from unmanaged populations (ß = -0.19, 95% CI = -0.24–-0.14, *P* = 6.43^e-9^, Figure 1A). This pattern was driven by both the number and length of ROH, the former being almost three times higher in the most inbred population than in the least inbred population (Figure 1B and Figure S5).

**Figure 1.**
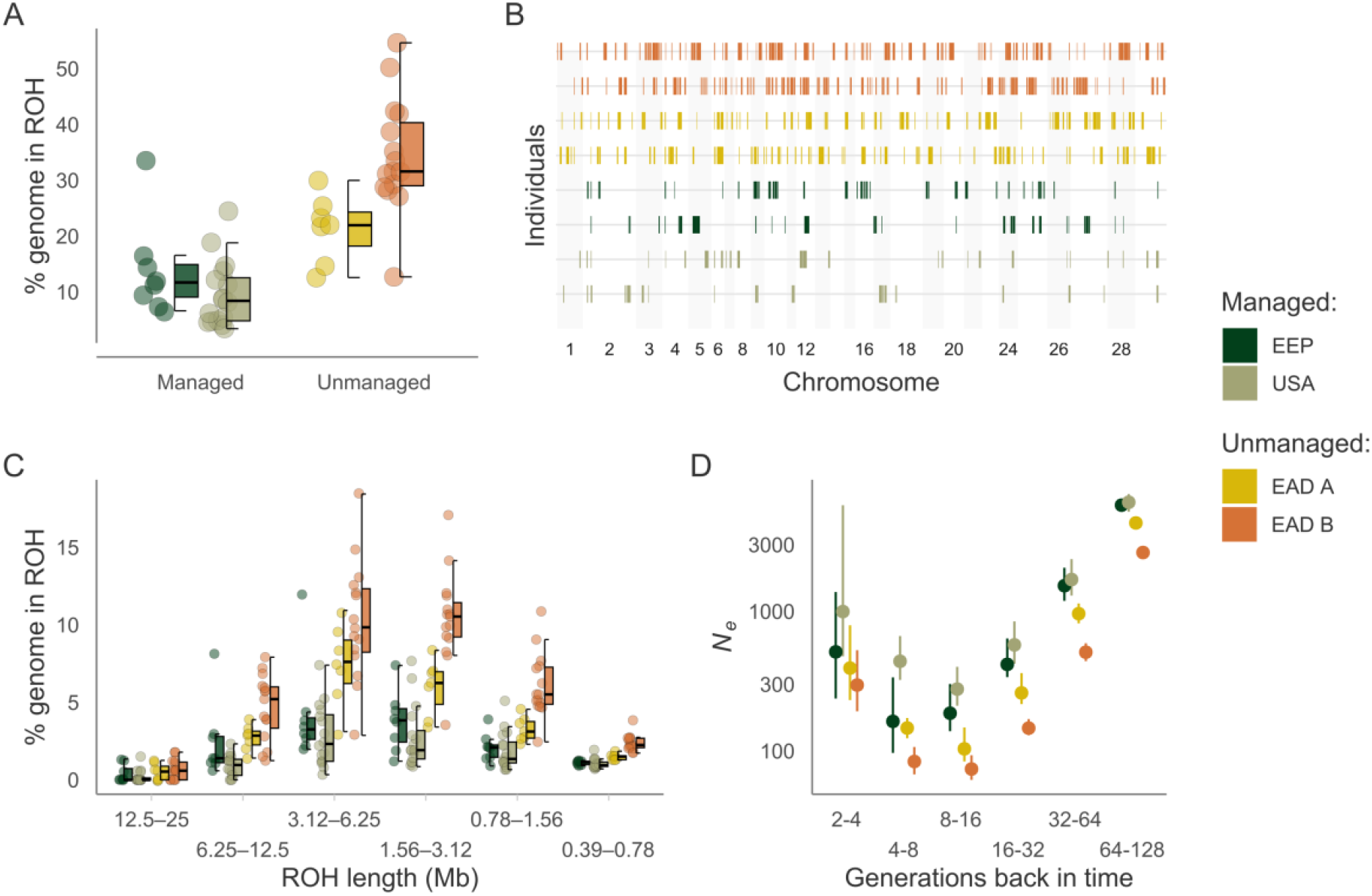
Runs of homozygosity (ROH) landscape across contrasting management strategies of scimitar-horned oryx. **(A)** Distribution of *F*_ROH_ across scimitar-horned oryx management strategies. Values were multiplied by 100 to reflect the percentage of the autosomal genome in ROH. Centre lines of boxplots reflect the median, bounds of the boxes extend from the first to the third quartile and upper and lower whiskers reflect variability outside the interquartile range. **(B)** ROH in the two individuals with intermediate inbreeding coefficients *F*_ROH_ from each population. **(C)** Distribution of ROH within different length classes. Data points represent the percentage of ROH of a given length within an individual’s autosomal genome. **(D)** Effective population size estimates inferred from the mean *F*_ROH_ in a population for a given time-period (see Materials and Methods for details). Error bars represent 95% bootstrap confidence intervals.

### ROH length distribution and recent demography

We also observed variation in the fraction of the genome in ROH of different length classes. There was a steep decrease in the genome fraction in ROH above lengths of around 6.25 Mb (Figure 1C). ROH longer than this made up a relatively small fraction of the genome, reaching a minimum average frequency of 0.4% between 12.5–25 Mb. ROH between 3.12–6.25 Mb had the highest frequency, making up on average 6.2% of an individual’s genome. Although this pattern was observed in each population, there was variation in absolute proportions across individuals. For example, length class 3.12–6.25 Mb made up on average only 3% of the genome in the least inbred population, USA, while it comprised on average 10% in the most inbred population, EAD B (Figure 1C). Interestingly, long ROH >12.5 Mb, which are likely the result of recent shared parental ancestry, were identified in less than 30% of individuals from managed populations, yet were present in over 60% of individuals from unmanaged populations.

As ROH lengths decrease, their underlying haplotypes are expected to have originated from ancestors further back in time (46). The genome fraction in ROH of different length classes can therefore provide insights into past changes in effective populations size (*N*_*e*_) (47, 48). In line with this, our estimates of *N*_*e*_ based on individual inbreeding coefficients were inversely proportional to the fraction of the genome in ROH of different length classes (Figure 1D). *N*_*e*_ declines to reach a minimum of around 150 individuals between 8–16 generations ago, after which it shows a steady increase towards the present day. Managed populations had higher *N*_*e*_ estimates across all time-periods than unmanaged populations (mean *N*_*e*_: USA = 1,672, EEP = 1,429 versus EAD A = 1,028, EAD B = 625, see also Figure S6). These patterns were reflected in estimates of mean pairwise nucleotide diversity which were also higher in managed (USA = 0.46 × 10^5^, EEP = 0.44 × 10^5^) than unmanaged populations (EAD A = 0.42 × 10^5^, EAD B = 0.27 × 10^5^).

### Mutation load landscape across management strategies

We next investigated how mutation load varies across management strategies using metrics based on putative deleterious variants identified using annotation-based methods. As the overall patterns were qualitatively similar across two variant effect prediction software (Figure S7), results using annotations from SnpEff are presented here. We first estimated two components of mutation load; heterozygous and homozygous mutation load for both weakly (missense) and highly (loss of function) deleterious mutations. The heterozygous mutation load was calculated as the absolute number of missense and loss of function heterozygotes per individual. The homozygous mutation load was calculated as the absolute number of derived missense and loss of function homozygotes per individual. Based on the assumption that most deleterious mutations are (at least) partially recessive (2, 49–51) and therefore partially hidden from selection when in a heterozygous state, we considered the homozygous mutation load to be most informative of the fitness cost due to inbreeding in the current population, and the heterozygous mutation load to reflect the potential for inbreeding to reduce fitness in future generations.

Heterozygous mutation load for both missense and loss of function (LoF) mutations was consistently higher in managed than unmanaged populations (Missense: ß = 1079, 95% CI = 769–1390, *P* = 1.12^e-8^, LoF: ß = 21.7, 95% CI = 12.7–30.6, *P* = 1.51^e-5^, Figure 2A–B). As expected, this pattern inversely tracked overall inbreeding levels, where individuals with lower inbreeding coefficients had a larger number of heterozygotes at missense and loss of function sites (Figure S8). In direct contrast, the homozygous mutation load for both missense and LoF mutations was lower in managed than in unmanaged populations (Missense: ß = -608, 95% CI = -779–-437, *P* = 6.41^e-9^, LoF: ß = -13.6, 95% CI = -18.3–-8.86, *P* = 6.99^e-7^, Figure 2C–D). Of the homozygous missense genotypes, 1,763 (11.5%), 6,790 (17.1%), 236 (0.81%) and 874 (5.43%) were due to alleles fixed in EAD A, EAD B, USA and EEP, respectively. Of the homozygous LoF genotypes, 173 (37.5%), 313 (28.4%), 46 (5.07%) and 95 (20.1%) were due to alleles fixed in EAD A, EAD B, USA and EEP, respectively.

**Figure 2.**
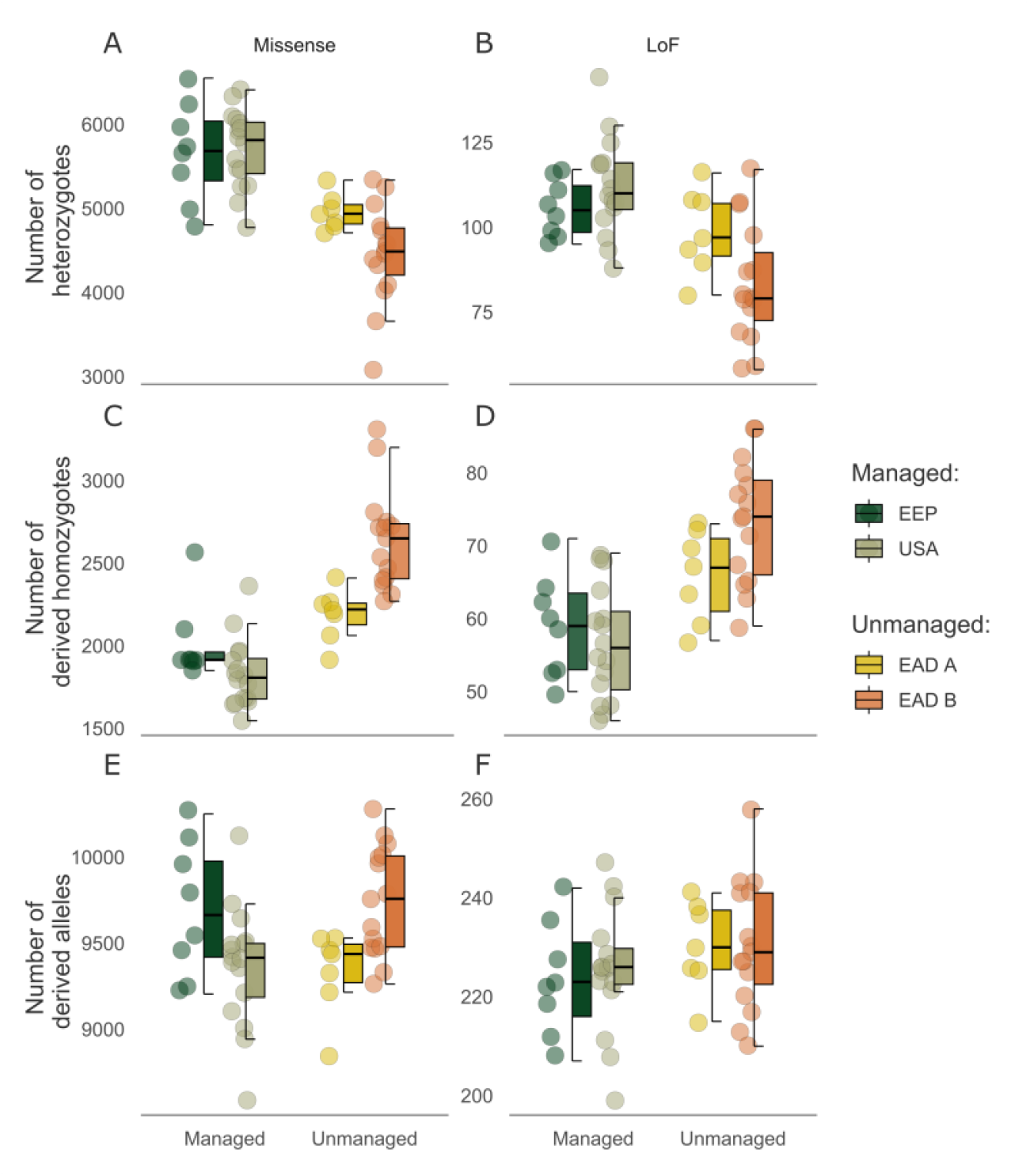
Deleterious load landscape across contrasting management strategies of scimitar-horned oryx based on SNPeff annotations. Distribution of the number of heterozygotes per individual (heterozygous mutation load) for missense **(A)** and loss of function mutations **(B)** across management strategies. Distribution of the number of derived homozygotes per individual (homozygous mutation load) for missense **(C)** and loss of function **(D)** mutations across management strategies. Distribution of the total number of derived alleles per individual for missense **(E)** and loss of function **(F)** mutations across management strategies.

To account for a scenario where deleterious mutations are approximately additive, we also calculated the total number of derived deleterious alleles per individual (52). Here, no significant difference in the total of number of LoF or missense alleles was observed between managed and unmanaged populations (Missense: ß = 136, 95% CI = -83.7–357, *P* = 0.22, LoF: ß = 5.53, 95% CI = -1.48–12.5, *P* = 0.12, Figure 2E–F). In addition, we used the measure *Rxy* to determine whether there was an excess of putative deleterious mutations in one management strategy over another. *Rxy* compares the relative frequency of derived alleles within a given impact category and is standardised over a set of intergenic SNPs, making it robust to population-specific biases. Unmanaged populations displayed a marginal excess of missense mutations compared to managed populations while no difference in the frequency of highly deleterious loss of function mutations could be detected between the management groups (Figure S9).

## Discussion

The scimitar-horned oryx was declared Extinct in the Wild in 2000, yet the species has persisted *ex-situ*. Understanding how management shapes the genomic landscape of inbreeding and mutation load is essential for supporting species viability. We used whole genome resequencing data to characterise runs of homozygosity and deleterious mutations in scimitar-horned oryx populations undergoing contrasting management strategies. Our study highlights the dynamics between inbreeding, mutation load and population size and has broad-reaching implications for practical conservation management.

We first demonstrated how signatures of recent population history can be identified in the genomes of present-day animals. Across *ex-situ* oryx populations, both managed and unmanaged, we observed a peak in the genome fraction in ROH between 3.12–6.25 Mb. Although it is not possible to precisely estimate the time to the most recent common ancestor (MRCA) when ROH are inferred using physical positions (53), the expected coalescent time for ROH of this size is approximately 8–16 generations ago (46). This shift in the genome fraction in ROH indicates a smaller population size around this time-period which we could reconstruct with our measures of *N*_*e*_. Interestingly, assuming a generation time of around seven years (44), this directly corresponds to the mid-20^th^ century when oryx were close to extinction in the wild and when *ex-situ* populations were founded (15, 43). We observed highly comparable patterns of *N*_*e*_ with the software GONe, together highlighting the power of ROH and LD-based methods for inferring the strength and timing of recent bottlenecks and for placing contemporary patterns of nucleotide diversity into a historical context.

The overall pattern of ROH lengths was qualitatively similar across populations yet the absolute proportion of the genome in ROH was considerably lower in managed than unmanaged populations for all length classes. Long ROH are indicative of recent shared ancestry of the underlying haplotypes because recombination has had little opportunity to break them up (54–57). The relative absence of long ROH therefore strongly indicates that close inbreeding is uncommon in managed populations, which work to mitigate this process. Furthermore, the smaller proportion of short ROH suggest managed populations also have lower levels of background relatedness (56, 58). Historic data on the origins of the unmanaged populations are lacking (15), yet it is not unreasonable to expect a higher level of relatedness among founder individuals compared to those of breeding programmes. Overall, these findings reveal the genomic effects of multiple generations of inbreeding, while on the other hand demonstrate how 30–40 years of *ex-situ* management has been successful at maximising the genome-wide diversity of captive populations.

We next shed light on the relationship between inbreeding, diversity and deleterious mutations by exploring how mutation load compares across management strategies. At an individual level, we show that animals from collections employing genetic management practices have a higher heterozygous mutation load for both missense and loss of function mutations than animals from unmanaged populations. Indeed, with lower levels of inbreeding – as observed in unmanaged SHO populations – a greater number of heterozygotes are expected. Furthermore, theory and simulations predict that large populations will have higher frequencies of segregating deleterious mutations (20, 59, 60). This is in part due to being masked from the effects of purifying selection in populations with larger *N*_*e*_, but also by genetic drift driving deleterious mutations to fixation in small populations. In line with this, we show that managed populations of oryx have higher nucleotide diversity and effective population sizes than unmanaged collections.

The presence of segregating deleterious alleles within insurance populations may be considered a concern for conservation management. Indeed, there has been recent debate surrounding the risks associated with sourcing individuals for restoration from large, genetically diverse populations, given the higher expected levels of strongly deleterious variation (18, 19). However, these concerns are primarily relevant for populations predicted to remain small and isolated with high levels of inbreeding. Restoration programs that follow established IUCN/SSC guidelines (39) will seek to source individuals from genetically differentiated populations, release large numbers of animals over extended time frames and maximise initial population growth rate. When *ex-situ* populations and reintroductions are managed in such a way, recessive deleterious mutations are more likely to remain partially masked as heterozygotes (16, 61–63), as we demonstrate is the case in the *ex-situ* population of oryx. Furthermore, high genetic diversity in reintroduced populations should theoretically enrich adaptive potential due to an increase in additive genetic variance (20, 64–66). The scimitar-horned oryx reintroduction has followed best practice guidelines having so far released over 250 animals over a five-year time-period, and in eight separate release batches. Consequently, the released population has now reached close to 400 individuals, with over 150 calves born in the wild. Follow-up monitoring of release herds will provide a rare opportunity to validate these efforts within the context of a large-scale reintroduction effort.

In addition to the heterozygous mutation load, we also considered how both the homozygous mutation load and the total number of derived deleterious alleles varied across populations. Several recent studies have demonstrated reductions in the relative number of highly deleterious mutations in small versus large (21–23, 26, 37) and in modern versus historical populations (24, 25), and attributed these differences to the effects of purifying selection. We observed no difference in the total number of derived loss of function alleles between unmanaged and managed populations, nor any deficit in the *Rxy* analysis. Instead, we find an increase in the homozygous mutation load for loss of function mutations in unmanaged collections. This suggests that while the total number of alleles has remained unchanged, inbreeding has increased the homozygosity of strongly deleterious variants. Consequently, unmanaged populations of oryx may carry a higher fitness cost associated with a greater burden of homozygous genotypes. In line with this, there is some anecdotal evidence of high disease prevalence and low juvenile survival in individuals originating from EAD B.

Similarly, missense mutations displayed no difference in total number between managed and unmanaged populations. However, higher values were observed in EAD B and EEP, the two populations thought to have experienced the strongest bottlenecks (see Supplementary Material). Missense mutations are more likely to be weakly deleterious with higher starting allele frequencies than loss of function mutations, and therefore are predicted to be more susceptible to accumulation by drift when population size is reduced (67–70). Critically however, the proportion of missense alleles that could be attributed to homozygous genotypes was higher in the unmanaged populations. If deleterious mutations are assumed to be approximately additive, the total number of derived alleles will provide a better estimate of genetic load and by extension be more informative of fitness (52, 71). On the other hand, if deleterious mutations are closer to recessive (2, 49–51), the homozygous mutation load will be more instructive. Indeed, inbreeding depression is ubiquitous in wild mammal populations (29, 33, 72–76), indicating that deleterious alleles have the most fitness impact when homozygous. This is exemplified by empirical work on a small, isolated population of Soay sheep with an estimated *N*_*e*_ of 194 that is thought to have remained in the low hundreds for thousands of years (77, 78). Despite this, long-term fitness and genomic data have revealed strong inbreeding depression caused by the expression of many weakly deleterious mutations (35, 72). With this in mind, even under a mutation load burden, *ex-situ* management to maximise genetic diversity will likely mask the effects of many deleterious alleles. Nevertheless, until we can validate molecular measures of load against empirical inbreeding depression, we must exercise caution when inferring fitness effects. Moreover, it is important to consider that even in the presence of reliable estimates of inbreeding depression, understanding the consequences on population viability is not trivial. This is because the demographic consequences of natural selection are dependent on the ecological context (33, 79–81) and therefore cannot be predicted with molecular data alone.

*Ex-situ* breeding and species reintroduction planning are ultimately exercises in risk management, with genetics making up only one component of a multifaceted set of considerations (39). Overall, our study provides empirical support for the value of genetic management not only for minimising inbreeding, but also for producing populations with enhanced genetic diversity for adaptation to changing environmental conditions and release back into the wild (7, 63, 82–85). Indeed, as part of the World Herd approach (86), mixing of animals from multiple collections is now a key part of the scimitar-horned oryx reintroduction strategy. While such actions can largely be informed using traditional measures of genetic variation, our study demonstrates how the application of whole genome sequencing in the context of *ex-situ* management has the power to resolve previously unknown aspects of variation. We recognise that it is impractical to consider comprehensive genomic approaches for the genetic management of every species (87). Rather, we suggest the application of studies such as this to guide conservation breeding strategies across diverse taxa and highlight the need for further work to link molecular predictions of inbreeding depression with empirical demographic data.

## Materials and methods

### Sampling and sequencing

Blood (in EDTA) and tissue (in 100% ethanol) samples were collected for whole genome resequencing from 49 scimitar-horned oryx representing four *ex-situ* populations: the EEP (*n* = 8), USA (*n* = 17), EAD A (*n* = 9) and EAD B (*n* = 15). The EEP and USA are captive collections undergoing genetic management practices, while EAD A and EAD B represent collections in the United Arab Emirates with no genetic management in place (see Supplementary Methods). Total genomic DNA was extracted between one and five times per sample using the DNeasy Blood and Tissue Kit (Qiagen, Cat. No. 69504). Elutions were pooled and concentrated in an Eppendorf Concentrator Plus at 45°C and 1400 rpm until roughly 50 µl remained. Library construction was carried out using the Illumina TruSeq Nano High Throughput library preparation kit (Illumina CA, UKA). Twenty samples from across all four populations were 150 bp paired-end sequenced on an Illumina HiSeq X Ten platform at a target depth of coverage of 15X. The remaining 29 samples from three of the populations were 150 bp paired-end sequenced on an Illumina NovaSeq 6000 instrument at a target depth of coverage of 7X (Table S1).

### Read processing and alignment

Sequence reads were assessed for quality using FastQC v0.11.7 (88) and trimmed for adaptor content using cutadapt v1.16 (89). Reads were then mapped to the scimitar-horned oryx reference genome assembly (*Oryx dammah* assembly version 1.1, Genbank accession number GCF_014754425.2) using BWA MEM v0.7.17 (90) with default parameters. Unmapped reads were removed from the alignment files using SAMtools v1.9 (91). Alignments were then sorted, read groups added and duplicates removed using Picard Tools v2.18.16. This resulted in a set of 49 filtered alignment files, one for each resequenced individual. To account for coverage variation in our data (92), we used SAMtools to downsample our 20 high coverage alignment files to approximately 6X, which was the average depth of coverage of our low coverage samples. All subsequent analyses were carried out on the set of alignments with comparable coverage.

### Variant calling and filtering

Haplotype Caller and GenotypeGVCFs in GATK v3.8 (93) were used for joint genotyping across all samples. The resulting SNP data were filtered for biallelic sites using BCFtools v1.9 (94). To obtain a high-quality set of variants we then used VCFtools (95) to remove loci with a quality score less than 30, a mean depth of coverage less than 5 or greater than 20, a genotyping rate less than 95% and a minor allele count less than 1. We removed SNPs originating from the X chromosome or any of the unplaced scaffolds within the assembly. One individual with a high relatedness score was dropped from subsequent analysis (Figure S10, see Supplementary Methods for details). The resulting SNP dataset contained over 10 million polymorphic sites with a genotyping rate of 98%.

### Population structure

We characterised population structure using genotype likelihood-based approaches in NGSadmix (96) and PCAngsd (97). Genotype likelihoods were first estimated from bam files in ANGSD (98) using the GATK model (-GL 2), inferring major and minor alleles (-doMajorMinor 1) and outputting only polymorphic sites (-SNP_pval 1e-6) with data in at least 60% of individuals (-minInd 30). We restricted this analysis to the 28 chromosome-length autosomes and included only regions with Phred quality and mapping scores over 30. Admixture proportions for the individuals in our dataset were calculated using NGSadmix. We performed admixture runs for ancestry clusters ranging from *K*=1–6, with ten runs for each *K*. The runs with the highest likelihood were plotted. The optimal *K* was identified based on the maximum value of the mean estimated *ln* probability of the data (99) and the Delta K method (100). Two individuals with intermediate admixture proportions between EAD A and EAD B were dropped from further analysis (Figure S2, see Supplementary Methods for details). We then performed a principal components analysis (PCA) using PCAngsd with the default parameters. Eigenvectors were computed from the covariance matrix using R.

### ROH calling and individual inbreeding coefficients

We used the filtered SNP genotypes to estimate inbreeding as the proportion of the genome in runs of homozygosoty (*F*_ROH_). ROH were called with a minimum length of 500 kb and a minimum of 50 SNPs using the --homozyg function in PLINK v1.9 (101) and the following parameters: --homozyg-window-snp 50 --homozyg-snp 50 --homozyg-kb 500 --homozyg-gap 1000 --homozyg-density 50 --homozyg-window-missing 5 --homozyg-window-het 3. We then calculated individual inbreeding coefficients *F*_ROH_ as the sum of the detected ROH lengths for each individual over the total autosomal assembly length (2.44 Gb). To ensure that our inbreeding estimates were not confounded by coverage, we compared *F*_ROH_ with the mean sequencing depth for each individual (Figure S11). Next, we ran linear models to explore the effect of management on inbreeding coefficients with *F*_ROH_ as the response variables and management strategy as the predictor variable. We also calculated *F*_ROH_ based on ROH inferred using bcftools roh and the following parameters: --AF-dflt 0.16 (average minor allele frequency), -G 30 and -M 1.2 (cattle recombination rate, Mouresan *et al*. 2019). We observed a near-perfect correlation (*r* = 0.99) with our PLINK-based estimates (Figure S12).

### ROH length distribution and recent demography

To assess recent changes in oryx population size, we characterised the fraction of the genome in ROH of seven different length classes (≥25, 12.5–25, 6.25–12.5, 3.12–6.25, 1.56–3.12, 0.78–1.56 and 0.39–0.78 Mb). Length classes (*L*) were calculated using the formula *L* = 100/(2 x g) (46), and reflect the expected lengths of ROH when the underlying haplotypes have most recent common ancestors <2, 2–4, 4–8, 8–16, 16–32, 32–64 and 64–128 generations (*g*) ago respectively. These generations were chosen to capture the time-period during which the wild population of oryx went extinct and captive populations were established. As there is no linkage map for the oryx, we assumed a genome-wide, homogenous, mean recombination rate of 1 cM/Mb and therefore used physical map lengths as opposed to genetic map lengths. For each time-period and for each individual, *F*_ROH_ was calculated as the sum of the ROH expected to coalesce in that time-period over the total autosomal assembly length (2.44 Gb). We then used measures of *F*_ROH_ to infer recent changes in effective population size across each population. For each time-period described above (*t)*, we first calculated the average *F*_ROH_ for each population. We then estimated *N*_*e*_ given the following expression where *F*_*ROH,t*_ corresponds to the average population inbreeding coefficient for a given time-period and *t* is the maximum number of generations in that time-period:

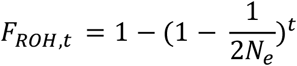

To calculate 95% bootstrap confidence intervals around our estimates, we resampled individuals within each population with replacement 100 times, and recalculated *N*_*e*_. In addition to this identity-by-descent (IBD) based approach, we implemented an LD-based method for estimating *N*_*e*_ using the software GONe (103). This is because under intensive inbreeding in recent generations, there are fewer ROH with long coalescent times making it more difficult to reliably estimate *N*_*e*_ in deeper history (see Supplementary Material for further information). As expected, historical estimates of *N*_*e*_ were larger using the LD-based approach, however the general pattern of population size change was the same across both methods (Figure S6).

### Nucleotide diversity

Nucleotide diversity estimates were calculated for each population using ANGSD. We first estimated the unfolded site-frequency spectrum (SFS) using the -doSaf and -realSFS commands while restricting the analysis to the 28 chromosome-length autosomes and regions with Phred quality and mapping scores over 30. Per-site pairwise nucleotide diversity estimates were then calculated using the -thetaStat command.

### Identification of deleterious mutations

As most deleterious mutations are likely to be derived alleles, we first polarised our SNP genotypes as ancestral or derived using the blue wildebeest (*Connochaetes taurinus*), topi (*Damaliscus lunatus*) and hartebeest (*Alcelaphus buselaphus*) as outgroup species. Short read sequencing data from wildebeest (SRR6902709), topi (SRR6913384) and hartebeest (SRR6922939 and SRR6922940) were downloaded from NCBI and mapped to the scimitar-horned oryx reference genome using BWA MEM with the default parameters. The alignments were then merged using SAMtools. A consensus was generated by selecting the most common base from the alignment using the doFasta 2 and doCounts 1 options in ANGSD. We then used PLINK v2.0 to polarise the oryx SNPs in our VCF based on the alleles in the consensus. First, we removed SNPs from our VCF whose positions were not present in the consensus sequence. Second, we removed SNPs where the ancestral allele in the consensus matched neither allele in the VCF file. Finally, we rotated alleles so that the reference allele in our VCF matched the ancestral allele in the consensus.

To identify deleterious mutations, we predicted the functional effects of the polarised SNP variants using both SnpEff v5.0 (104) and the Variant Effect Predictor (VEP) v99.2 (105). These methods compare a set of variants to an annotation database and predict the consequence of the derived alleles on genes, transcripts and proteins. Both were run using the NCBI RefSeq scimitar-horned oryx genome annotation downloaded from: https://ftp.ncbi.nlm.nih.gov/genomes/all/annotation_releases/59534/100/GCF_014754425.2_SCBI_Odam_1.1/. For each approach, sites with warnings were removed from the VCF file and SNPs were categorised as loss of function or missense according to the classifications provided in Table S2. 643 loss of function and 34,387 missense sites were identified with SnpEff and 760 loss of function and 42,440 missense were identified by the VEP. For each dataset, we also extracted a random subset of 100,000 intergenic SNPs. For each set of SNPs, genotypes were extracted for all individuals using a combination of VCFtools and PLINK.

### Mutation load landscape across management strategies

To assess how the mutation load varies across populations we used multiple measures. First, we approximated two components of mutation load; heterozygous and homozygous mutation load. The heterozygous mutation load was measured as the total number of heterozygotes per individual for both loss of function and missense variants. The homozygous mutation load was measured as the total number of derived homozygotes per individual for both loss of function and missense variants. Next, we calculated the total number of derived alleles per individual for both missense and loss of function variants. To ensure comparisons across groups were not confounded by patterns of neutral variation, we also calculated the total number of derived alleles at a random subset of 100,000 intergenic SNPs. No significant difference between management strategies was observed (ß = 139, 95% CI = -217–495, *P* = 0.44, Figure S13). Finally, we used the *Rxy* statistic to estimate the relative frequency of loss of function and missense mutations in one population over another (106). Derived allele frequencies were calculated based on individuals from managed and unmanaged populations separately. A random subset of 100,000 intergenic SNPs was used to standardise our estimates and account for population-specific biases. To calculate 95% bootstrap confidence intervals around our estimates, we randomly resampled SNPs with replacement 100 times, and recalculated *Rxy*. To explore the effect of management on genetic load, we ran linear models with homozygous mutation load, heterozygous mutation load or the total number of derived alleles as the response variable and management strategy as the predictor variable.

## Data availability

EEP samples are archived at the EAZA Biobank https://www.eaza.net/conservation/research/eaza-biobank. Whole genome resequencing data have been deposited to the European Nucleotide Archive under study accession number PRJEB37295. Analysis code is available at https://github.com/elhumble/SHO_roh_load_2022 and https://github.com/elhumble/SHO_reseq_2022.

## Author contributions

EH, RO, HS, ADB, and KD conceived and designed the study. JC, RP and MAR contributed materials and funding. BP provided samples from SSP populations. EH analysed the data with input from MAS and RMG. EH wrote the manuscript. All authors commented on and helped improve the final manuscript.

## Acknowledgements

We thank the EAD, EAZA and AZA institutions, along with private ranch owners in the USA, that provided samples for this study. We thank Jennifer Kaden for initial processing of samples and DNA extraction, and Edinburgh Genomics for carrying out the whole genome sequencing. SaharaConservation provided materials and wider project support. We also acknowledge Katerina Guschanski for helpful discussions and Tania Gilbert at Marwell Wildlife for information on the international studbook of scimitar-horned oryx. For the purpose of open access, the author has applied a Creative Commons Attribution (CC BY) licence to any Author Accepted Manuscript version arising from this submission.

