## Supplementary Material for "Conservation management strategy impacts inbreeding and mutation load in scimitar-horned oryx"

#### Table of contents

Supplementary Methods **Page 2**
Supplementary Tables **Page 6**
Supplementary Figures **Page 9**

### Supplementary Methods

**Population origins**

Scimitar-horned oryx have been kept in captivity since the 1800s yet the majority of known founders originated from a capture operation in Chad in the 1960s (1). As part of this, approximately 17 individuals were sent to Europe to form the basis of what is now the European Association of Zoos and Aquaria (EAZA) *Ex Situ* Programmes (EEP), and around 29 individuals were sent to the USA to form the basis of the Species Survival Plan® of the Association of Zoos and Aquariums (AZA) and privately-owned ranch populations (1), hereby referred to as the USA population. Large numbers of oryx are also housed in collections in the United Arab Emirates yet the founder origins of these populations are not well documented. Anecdotal evidence suggests individuals may have originated from across different historical range states (1).

Within the EEP, individuals undergo high-intensity genetic management to facilitate demographic stability and minimise inbreeding (2). Under this strategy, individual mean kinship and inbreeding coefficients derived from pedigree data are used to make breeding and transfer decisions. This approach should theoretically retain greater genetic diversity than under random mating (3). Typically, animals are kept in small herds and males are moved between collections to mimic natural dispersal patterns. No movement occurs with institutions outside the EEP. The current census size of the EEP population is around 619 individuals.

Within the USA there is a broader range of genetic management strategies in place. For example, some institutions within the SSP employ similar high-intensity genetic management practices as the EEP. However, this is not possible for all due to a lack of pedigree records and greater numbers of animals present in large enclosures (4). Under these scenarios, low-intensity genetic management takes place, where herds are assembled from multiple sources and bulls are regularly rotated among them. The current census size of the SSP population is around 223 individuals. In the USA, there are also substantial collections of scimitar-horned oryx held on privately-owned ranches (several thousand individuals). The management spectrum of ranch populations varies from low-intensity genetic management to collections of completely unmanaged breeding herds. Recently, increasing emphasis has been placed on metapopulation management within these private collections through the Source Population Alliance under the Conservation Centers for Species Survival initiative (5).

The Environment Agency – Abu Dhabi (EAD) also houses a population of several thousand scimitar-horned oryx in the United Arab Emirates. Most these animals were sourced from the late Sheikh Zayed bin Sultan Nahyan's private collection on Sir Bani Yas island (EAD A) which comprised thousands of individuals. An additional source came from a small population of around 70 animals that was identified in the UAE around 2014 (EAD B). Sparse historical records mean that founder numbers and origins for these source populations are unknown. However, prior to moving under the EAD's management purview, both were completely unmanaged and therefore provide a unique comparison group against managed EEP and USA populations.

**Sample selection**

Samples from the EEP population ($n$ = 8) originated from six European EAZA institutions employing high-intensity genetic management. Samples from the USA ($n$ = 17) originated from both private ranches (USA Ranch: $n$ = 10) and three AZA institutions employing a combination of high and low-intensity genetic management (USA AZA: $n$ = 7). Samples were selected to be genetically representative of the founding lineages within the EEP and USA populations. This was carried out on the basis of mtDNA control region haplotypes and was a necessary precaution as the genetically managed collections cannot be considered randomly mating populations. As we did not set out to carry out a systematic comparison of management strategies in the USA, we treated the USA Ranch and USA SSP samples as one population (USA: $n$ = 17) on the basis of the following:

(i) The ranch populations in which the samples originate are unknown, however anecdotal evidence suggest they are comprised of excess SSP animals and that, at a minimum, low-intensity genetic management is employed through regular bull rotation.

(ii) Analysis of population structure clustered USA SSP and USA Ranch individuals together with respect to the other sampled populations (Figure S1–3).

(iii) We identified one full-sibling pair in our dataset whereby one individual originated from a private ranch and the other from an AZA institution, corroborating the above.

(iv) We observe no differences in the main results of our manuscript between USA Ranch and USA SSP animals (Figures S14–15). Systematic sampling across a broader range of ranch types would provide greater power to investigate this in further detail.

In the EAD, samples were selected using origin information i.e. whether they originated from EAD source A or EAD source B. Due to the absence of management in these populations, we considered them to be representative of panmictic populations and therefore sampled individuals opportunistically. As the two source populations had been moved together prior to when sampling took place, we evaluated individual admixture proportions to ensure our samples were truly representative of the source populations (see Main Text for Methods). Two individuals were identified with intermediate ancestry between EAD A and EAD B and were removed from subsequent analysis (Figure S2).

**Relatedness**

To estimate relatedness among individuals, we first pruned our SNP dataset for linkage disequilibrium using the --indep function in PLINK, a sliding window of 50 SNPs, a step size of 5 and a variance inflation factor threshold of 2. We also removed SNPs that deviated significantly from HWE with a p-value threshold of 0.001 and with a minor allele frequency < 0.3. We then estimated KING, R0 and R1 coefficients (6) using NgsRelate v2 (7). These statistics are based on genome-wide patterns of identity by state sharing between two individuals. One pair of individuals (MSH648 / MSH682) fell above the KING-robust kinship threshold for first-degree relatives (Figure S10) (6) and therefore one individual from this pairing (MSH682) was removed from subsequent analysis. The remaining individuals displayed no sign of close relatedness (mean pairwise relatedness = $4.17 \times 10^{-3}$).

**Effective population size**

In addition to estimating effective population size ($N_e$) based on $F_{\text{ROH}}$, we also employed a linkage disequilibrium (LD) based approach for comparison. This is because under intensive inbreeding in recent generations, a large fraction of the genome is made up of long ROH with short coalescent times. As a result, there is less genomic territory for reliably estimating $N_e$ in deeper history, as demonstrated in Figure 2C. Furthermore, the performance and biases of IBD-based methods for estimating $N_e$ are generally not well understood. LD-based approaches such as GONe (8), use the observed patterns of linkage disequilibrium across chromosomes to estimate $N_e$ in the recent past. The approach can be applied to contemporary samples of fewer than 10 individuals making it particularly appropriate for our dataset.

The recommended number of SNPs per chromosome for analysis with GONe is between 50,000 and 100,000. We therefore randomly subsampled 2,000,000 filtered loci from across all 28 autosomes to achieve approximately ~70,000 SNPs per autosome. We then split the dataset into populations and ran GONe using the default parameters. The resulting $N_e$ trajectories showed a similar pattern across all populations, with a steep decline in $N_e$ to less than 100 individuals between 8 and 16 generations ago, corresponding to the time-period in which captive populations were founded (Figure S6). Furthermore, unmanaged populations (EAD A and EAD B) display lower $N_e$ estimates than managed populations (USA and EEP) across much of recent history. These patterns are highly comparable to those observed using the IBD-based approach. However, as expected, $N_e$ estimates in deeper history (>32 generations ago) were markedly larger using the LD-based method, at around 10-30K individuals. These values are likely to reflect more reliable estimates of historical $N_{e.}$ than those inferred using patterns of IBD. Indeed, the long-term $N_e$ estimate calculated for scimitar-horned oryx in (9) was around 22K individuals. Overall, these findings highlight the value of employing multiple methods for estimating $N_e$, particularly in populations with close inbreeding due to recent ancestry.

### Supplementary Tables

**Table S1.** International studbook #, sample ID, sex, origin, sample type, sequencing platform and sequencing coverage. EAD = Environment Agency Abu Dhabi, EEP = EAZA *Ex Situ* Programmes and SSP = American Association of Zoos and Aquariums Species Survival Plan®. NB: Possession of an international studbook ID does not necessarily mean an individual is managed through individual-based pedigree information.

| International studbook # | Sample ID | Sex | Origin | Sample type | Sequencing depth |
|---|---|---|---|---|---|
| NA | MSH753 | F | EAD A | Tissue | High |
| NA | MSH754 | F | EAD A | Tissue | High |
| NA | MSH755 | F | EAD A | Tissue | High |
| NA | MSH756 | M | EAD B | Tissue | High |
| NA | MSH757 | M | EAD B | Tissue | High |
| 34260 | MSH001 | M | EEP | Blood | High |
| 34412 | MSH005 | F | EEP | Tissue | High |
| 35552 | MSH009 | F | EEP | Blood | High |
| 30988 | MSH045 | M | EEP | Blood | High |
| 36828 | MSH054 | M | EEP | Blood | High |
| NA | MSH233 | M | EAD A | Blood | High |
| NA | MSH238 | F | EAD A | Blood | High |
| NA | MSH241 | M | EAD A | Blood | High |
| NA | MSH244 | M | EAD A | Blood | High |
| NA | MSH250 | F | EAD A | Blood | High |
| NA | MSH306 | M | EAD A | Blood | High |
| 33556 | MSH638 | NA | USA SSP | Blood | High |
| 37099 | MSH639 | NA | USA SSP | Blood | High |
| 36948 | MSH645 | NA | USA SSP | Blood | High |
| 37148 | MSH648 | NA | USA SSP | Blood | High |
| 39422 | MSH678 | F | USA Ranch | Blood | Low |
| 38070 | MSH682 | F | USA Ranch | Blood | Low |
| 39454 | MSH686 | F | USA Ranch | Blood | Low |
| 39318 | MSH687 | F | USA Ranch | Blood | Low |
| 39314 | MSH688 | F | USA Ranch | Blood | Low |
| 39459 | MSH693 | F | USA Ranch | Blood | Low |
| 39451 | MSH694 | M | USA Ranch | Blood | Low |
| 39456 | MSH695 | M | USA Ranch | Blood | Low |
| 39452 | MSH697 | M | USA Ranch | Blood | Low |
| 39319 | MSH699 | M | USA Ranch | Blood | Low |
| 39433 | MSH701 | M | EAD B | Blood | Low |
| 39504 | MSH703 | F | EAD B | Blood | Low |
| 38493 | MSH704 | F | EAD B | Blood | Low |
| 37603 | MSH705 | F | EAD B | Blood | Low |
| 38235 | MSH708 | M | EAD B | Blood | Low |

| | | | | | |
|---|---|---|---|---|---|
| 38681 | MSH710 | M | EAD B | Blood | Low |
| 38958 | MSH712 | M | EAD B | Blood | Low |
| 36871 | MSH725 | M | EAD B | Blood | Low |
| 37611 | MSH728 | M | EAD B | Blood | Low |
| 38236 | MSH731 | F | EAD B | Blood | Low |
| 41012 | MSH735 | M | EAD B | Blood | Low |
| 39507 | MSH737 | F | EAD B | Blood | Low |
| 38237 | MSH746 | F | EAD B | Blood | Low |
| 36932 | MSH641 | NA | USA SSP | Blood | Low |
| 36274 | MSH620 | NA | USA SSP | Blood | Low |
| 34176 | MSH656 | NA | USA SSP | Blood | Low |
| 37876 | MSH404 | F | EEP | Blood | Low |
| 37540 | MSH410 | M | EEP | Blood | Low |
| 36724 | MSH195 | M | EEP | Blood | Low |

**Table S2.** Classifications used to assign missense, loss of function and intergenic categories to variants annotated by SNPeff and VEP.

| Impact class | SNPeff | VEP |
|---|---|---|
| Missense | "missense_variant" | "missense_variant" |
| Loss of function | "LOF" | "transcript_ablation", "transcript_ablation", "splice_donor_variant", "splice_acceptor_variant", "stop_gained", "frameshift_variant", "inframe_insertion", "inframe_deletion", "splice_region_variant" |
| Intergenic | "intergenic_region" | "intergenic_variant" |

### Supplementary Figures

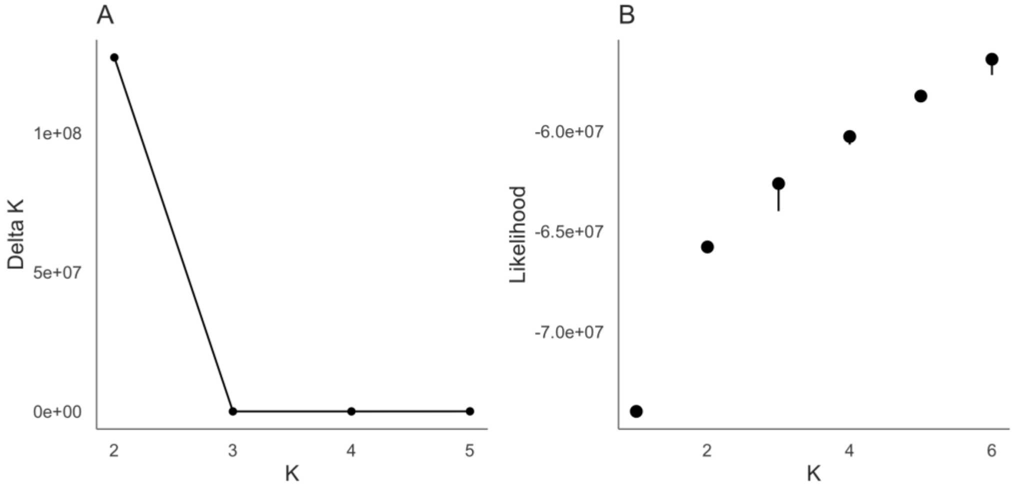


**Figure S1. (A)** Rate of likelihood change (Delta K) values based on ten replicate runs of NGSadmix for *K* = 2 to *K* = 6. **(B)** Mean log likelihood values calculated from ten replicate runs of NGSadmix for *K* = 1 to *K* = 6. Error bars reflect minimum and maximum values.

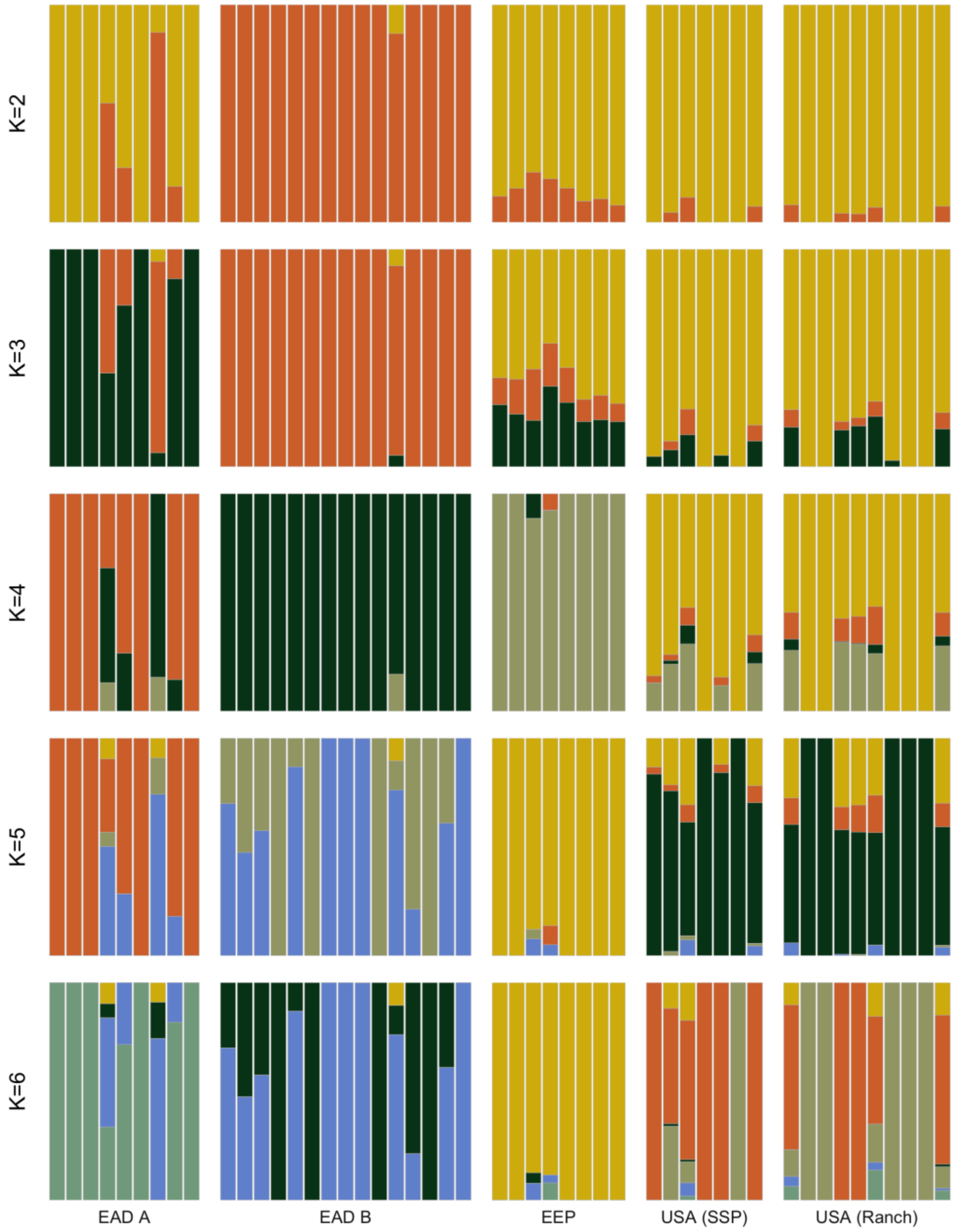


**Figure S2.** Ancestry proportions for each of the 49 scimitar-horned oryx individuals inferred using NGSadmix for $K$ = 2 to $K$ = 6.

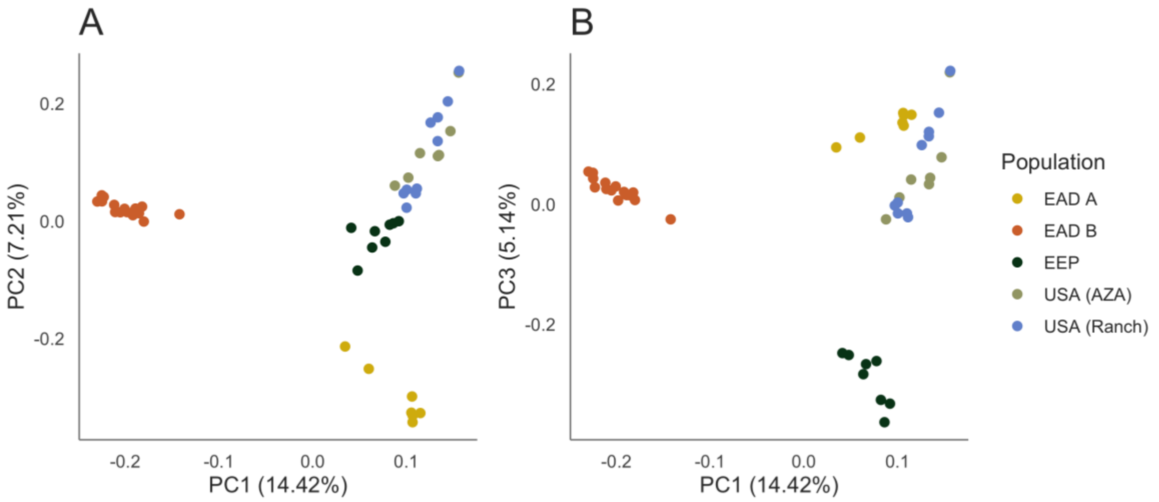


**Figure S3.** Scatterplots showing individual variation in principal components (PCs) one and two **(A)** and one and three **(B)** derived from principal components analysis in PCAngsd for 46 individuals. The amount of variance explained by each PC is shown in parentheses.

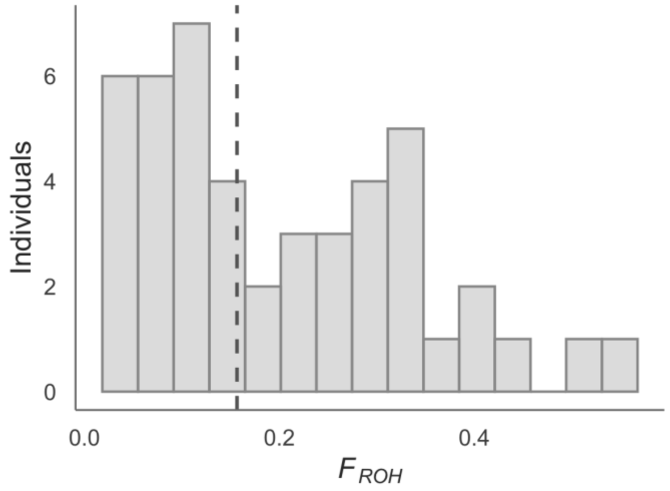


**Figure S4.** Distribution of inbreeding coefficients $F_{ROH}$ for 46 scimitar-horned oryx individuals. The dashed line represents the median.

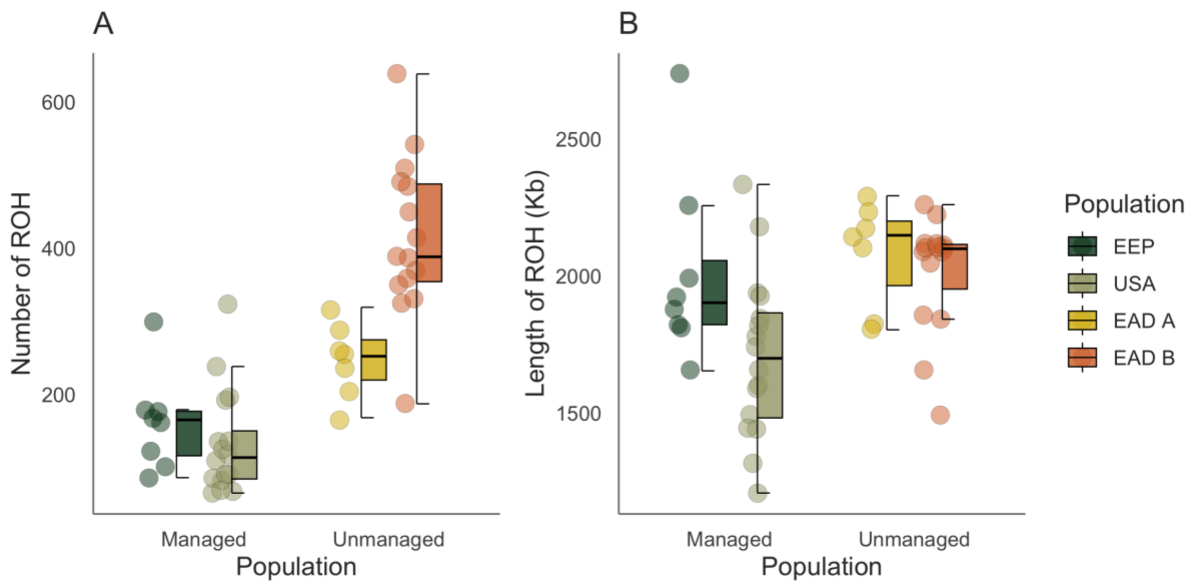


**Figure S5.** Distribution of the number **(A)** and length of ROH **(B)** across scimitar-horned oryx management strategies. Centre lines of boxplots reflect the median, bounds of the boxes extend from the first to the third quartile and upper and lower whiskers reflect variability outside the interquartile range.

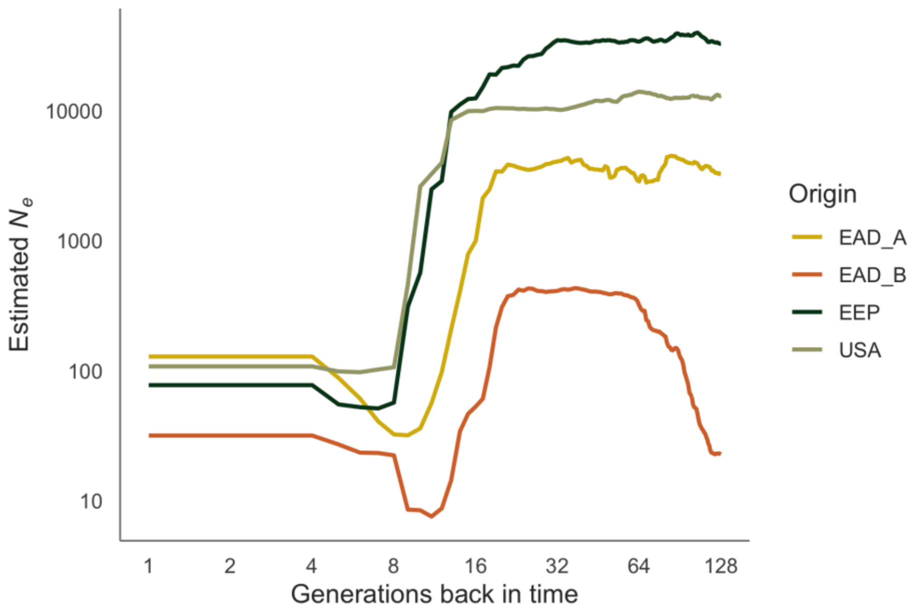


**Figure S6.** Effective population size estimates through time inferred using GONe for each population.

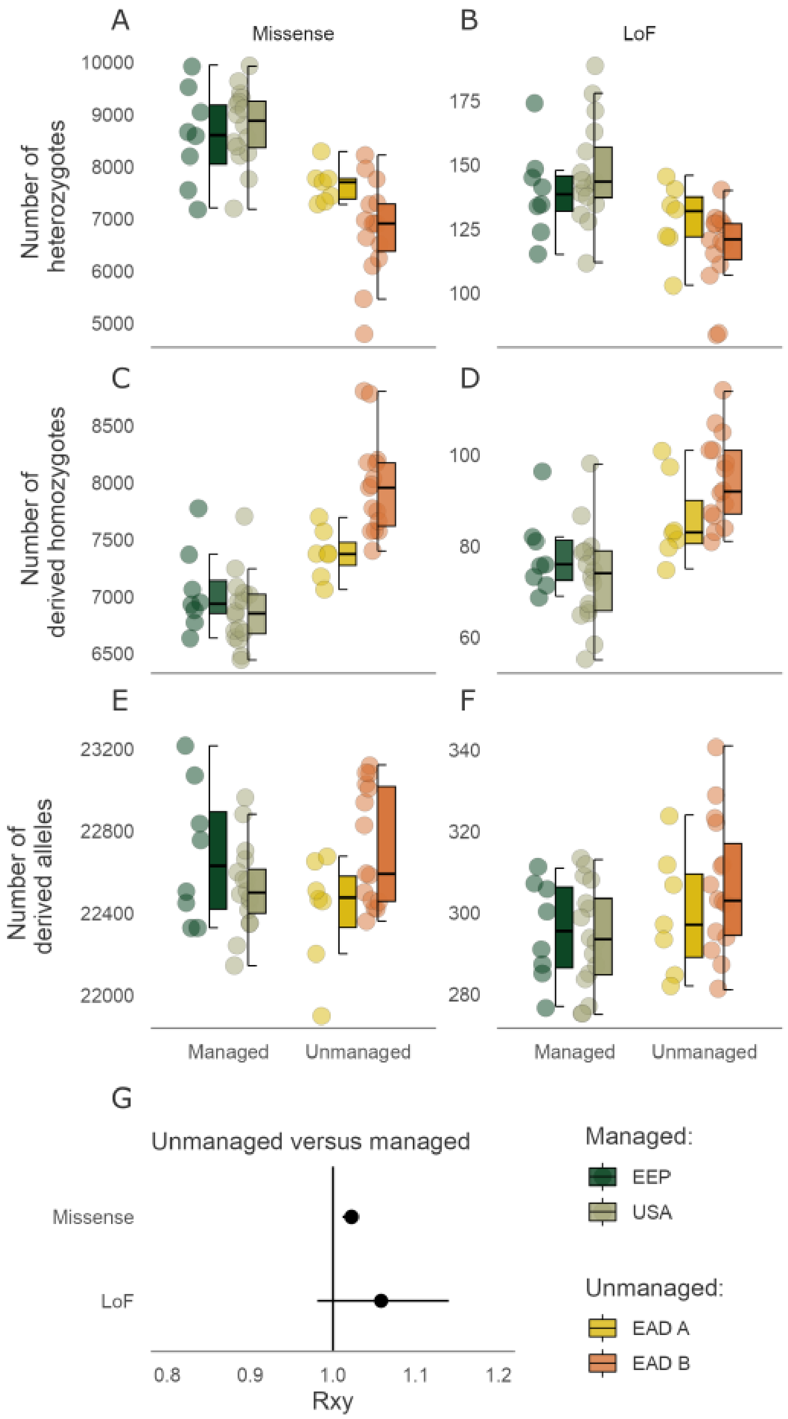


**Figure S7. Deleterious load landscape across contrasting management strategies of scimitar-horned oryx based on VEP annotations.** Distribution of the number of heterozygotes per individual (heterozygous mutation load) for missense **(A)** and loss of function mutations **(B)** across management strategies. Distribution of the number of alternative homozygotes per individual (homozygous mutation load) for missense **(C)** and loss of function **(D)** mutations across management strategies. **(D)** Distribution of the total number of derived alleles per individual for missense **(E)** and loss of function **(F)** mutations across management strategies. **(G)** Relative number (*Rxy*) of alternative alleles at missense and loss of function sites. *Rxy* > 1 indicates a relative frequency excess of a given category of sites in unmanaged versus managed populations. Error bars represent 95% bootstrap confidence intervals.

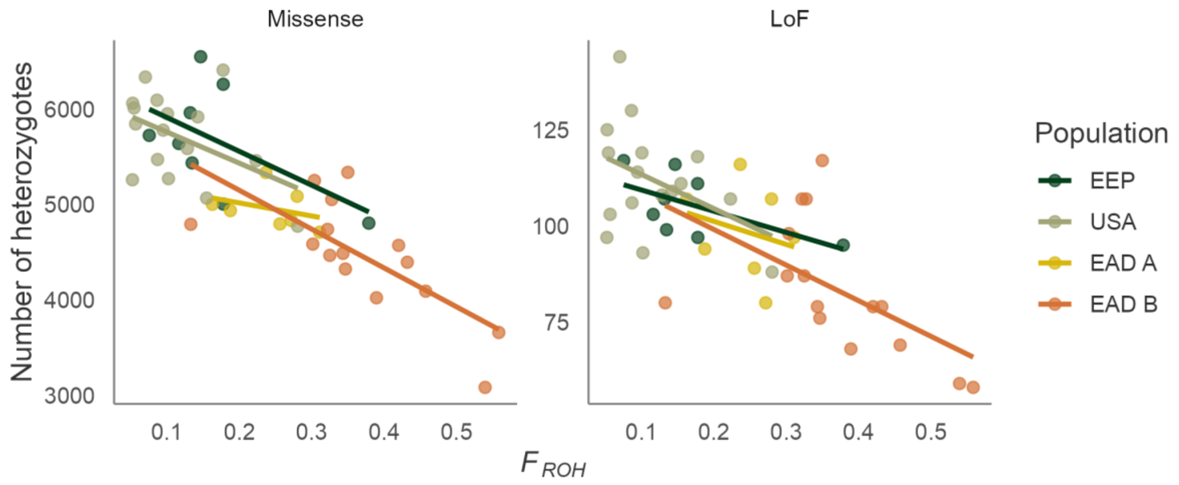


**Figure S8.** The heterozygous mutation load for missense and loss of function mutations plotted against the inbreeding coefficient $F_{ROH}$. Linear model regression lines shown for each population.

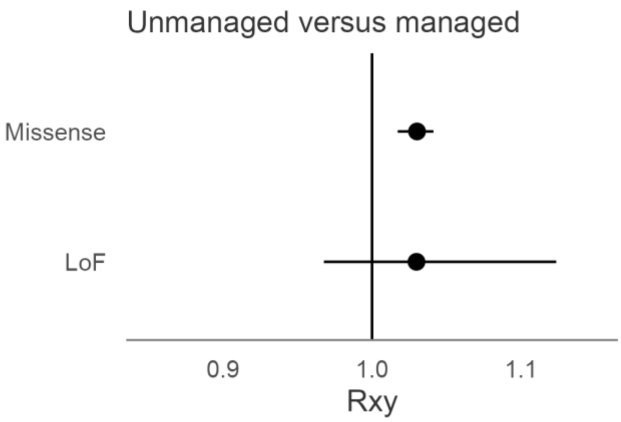


**Figure S9.** Relative number (*Rxy*) of derived alleles at missense at loss of function sites. *Rxy* > 1 indicates a relative frequency excess of a given category of sites in unmanaged versus managed populations. Error bars represent 95% bootstrap confidence intervals.

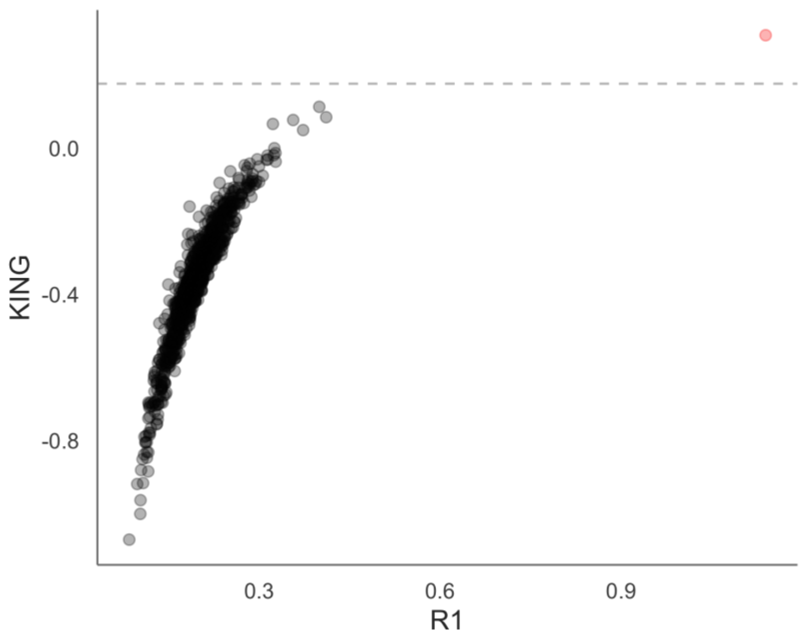


**Figure S10.** Empirical relatedness for all individual pairwise comparisons. R1 coefficients are plotted against KING-robust kinship coefficients. The dashed line represents the KING-robust kinship threshold statistic for first degree relatives and is equivalent to $\frac{1}{2\hat{}(5/2)}$ (6). One pair of individuals (MSH648 / MSH682) fell above this threshold and is colour coded in red.

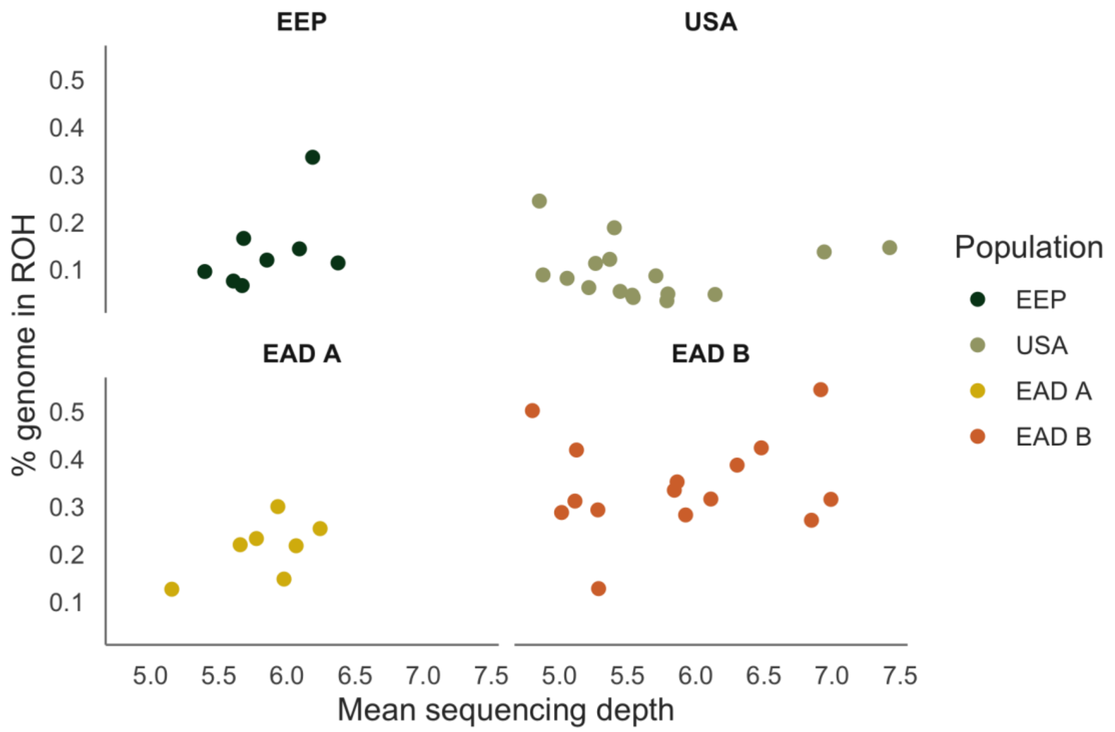


**Figure S11.** Mean sequencing depth plotted against the inbreeding coefficient $F_{ROH}$ for 46 individuals.

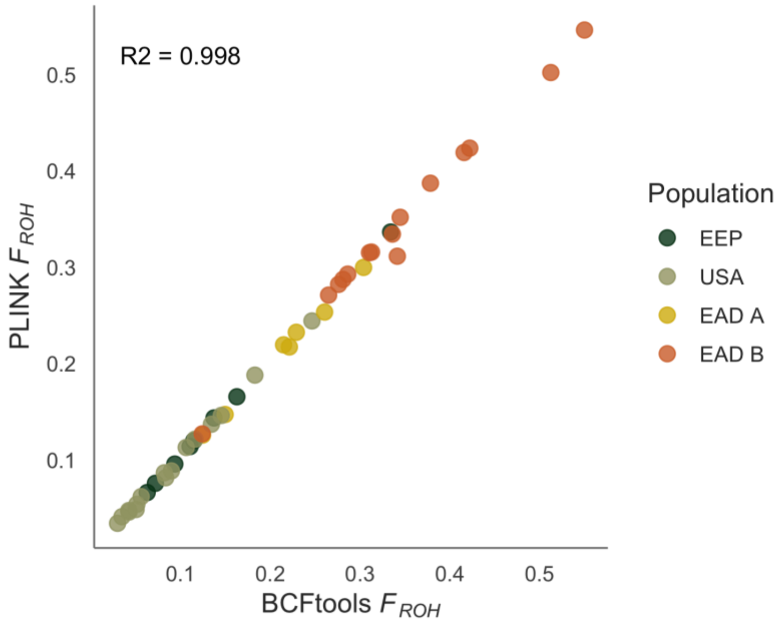


**Figure S12.** Pairwise correlation between inbreeding coefficients $F_{\mathrm{ROH}}$ based on ROH inferred using PLINK and BCFtools.

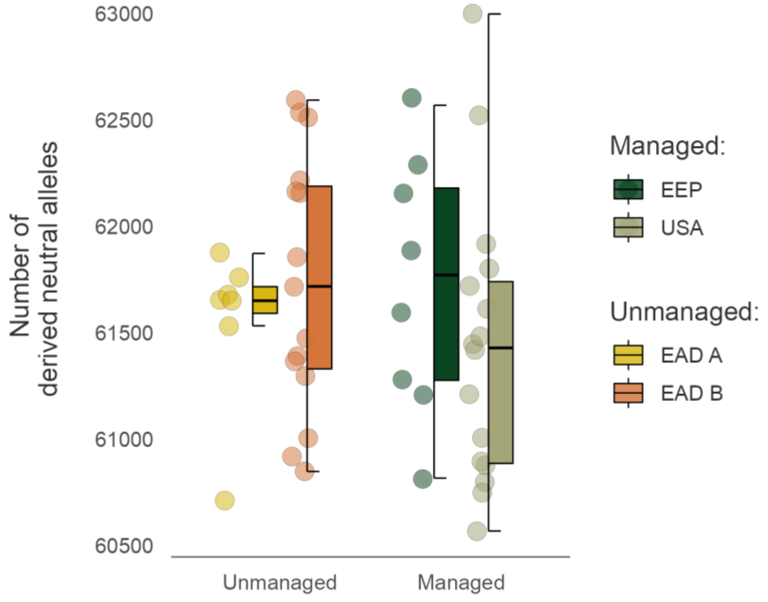


**Figure S13.** Distribution of the total number of derived alleles per individual for a random set of 100,000 intergenic (neutral) SNPs across management strategies.

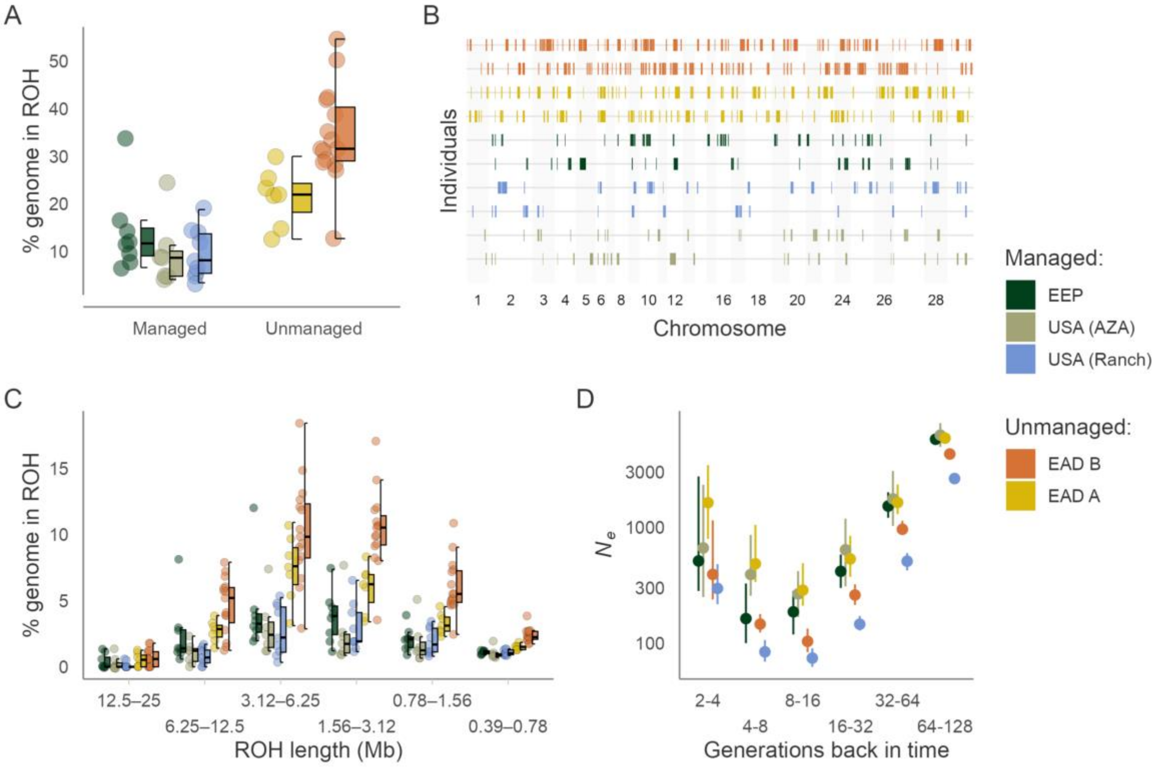


**Figure S14. Runs of homozygosity (ROH) landscape across contrasting management strategies of scimitar-horned oryx. USA (Ranch) and USA (AZA) animals are treated separately. (A)** Distribution of $F_{ROH}$ across scimitar-horned oryx management strategies. Values were multiplied by 100 to reflect the percentage of the autosomal genome in ROH. Centre lines of boxplots reflect the median, bounds of the boxes reflect the 25$^{th}$ and 75$^{th}$ percentile and upper and lower whiskers reflect the largest and smallest values. **(B)** ROH in the two individuals with intermediate inbreeding coefficients $F_{ROH}$ from each population. **(C)** Distribution of ROH within different length classes. Data points represent the percentage of ROH of a given length within an individual's autosomal genome. **(D)** Effective population size estimates inferred from the mean $F_{ROH}$ in a population for a given time-period (see Materials and Methods for details). Error bars represent 95% bootstrap confidence intervals.

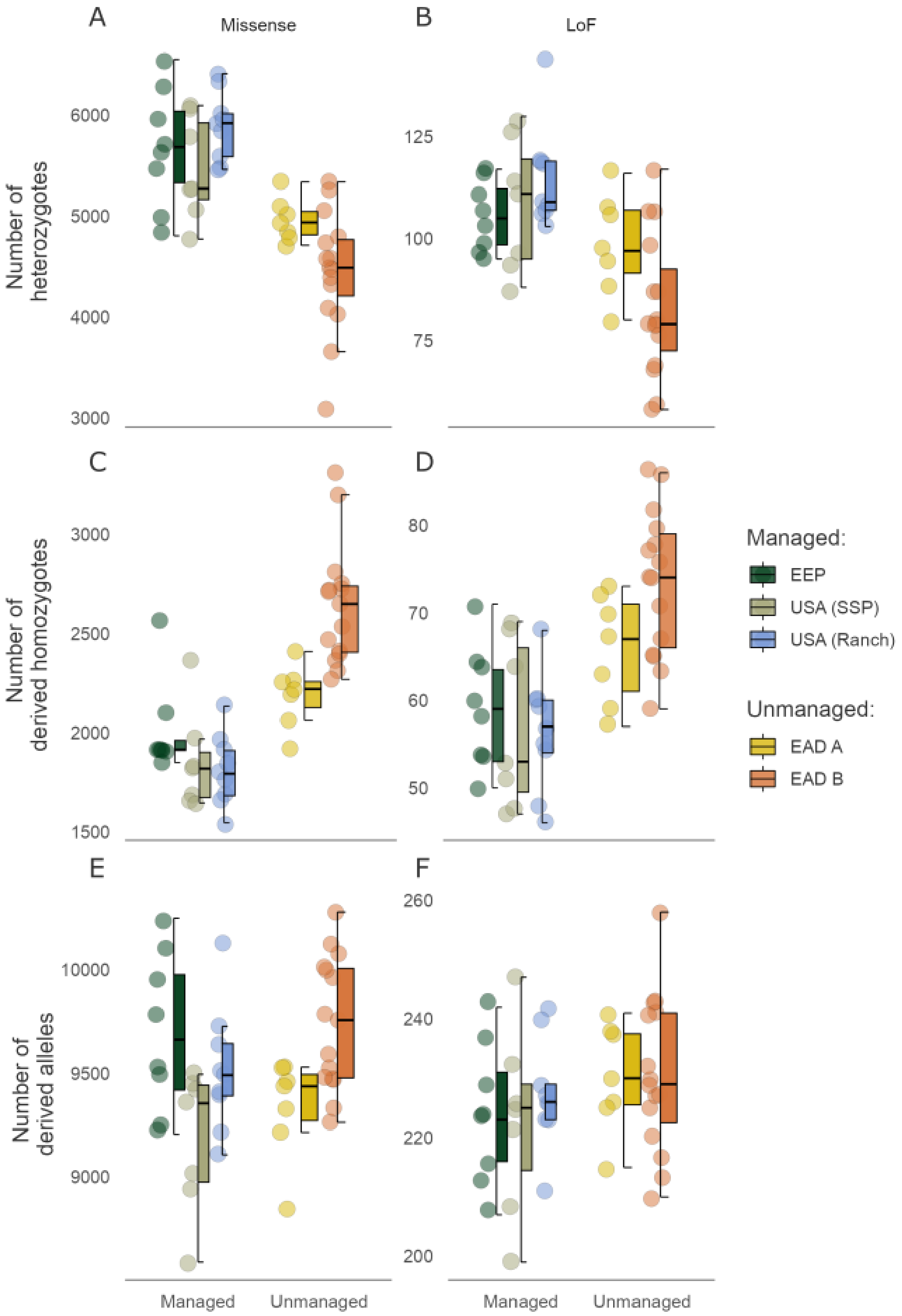


**Figure S15. Deleterious load landscape across contrasting management strategies of scimitar-horned oryx based on SNPeff annotations. USA (Ranch) and USA (AZA) animals are treated separately.** Distribution of the number of heterozygotes per individual (heterozygous mutation load) for missense **(A)** and loss of function mutations **(B)** across management strategies. Distribution of the number of alternative homozygotes per individual (homozygous mutation load) for missense **(C)** and loss of function **(D)** mutations across management strategies. Distribution of the total number of derived alleles per individual for missense **(E)** and loss of function **(F)** mutations across management strategies.
